## Supplementary Figures for "Biochemical-free enrichment or depletion of RNA classes in real-time during direct RNA sequencing with RISER"

### Supplementary Information

#### Supplementary Figures

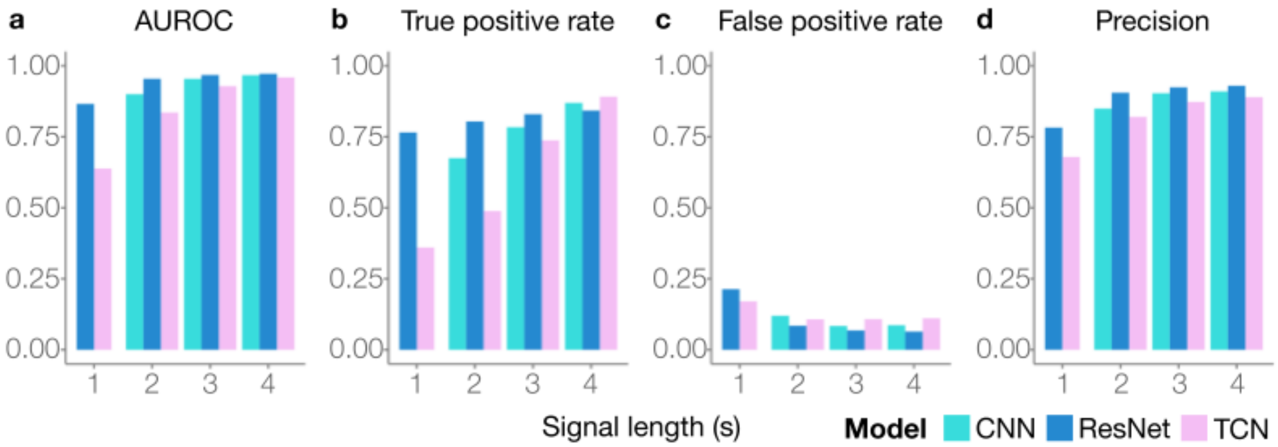


**Supplementary Figure 1: Evaluation of candidate RISER models**. **a-d,** Model performance on the test set for each candidate input signal length (x-axes) in seconds, color-coded by three convolutional network architectures: ”vanilla” convolutional neural network (CNN) (cyan), residual network (ResNet) (dark blue) and temporal convolutional network (TCN) (pink). We show the area under the receiver operating characteristic curve (AUROC) (**a**), true positive rate (**b**), false positive rate (**c**) and precision (**d**).

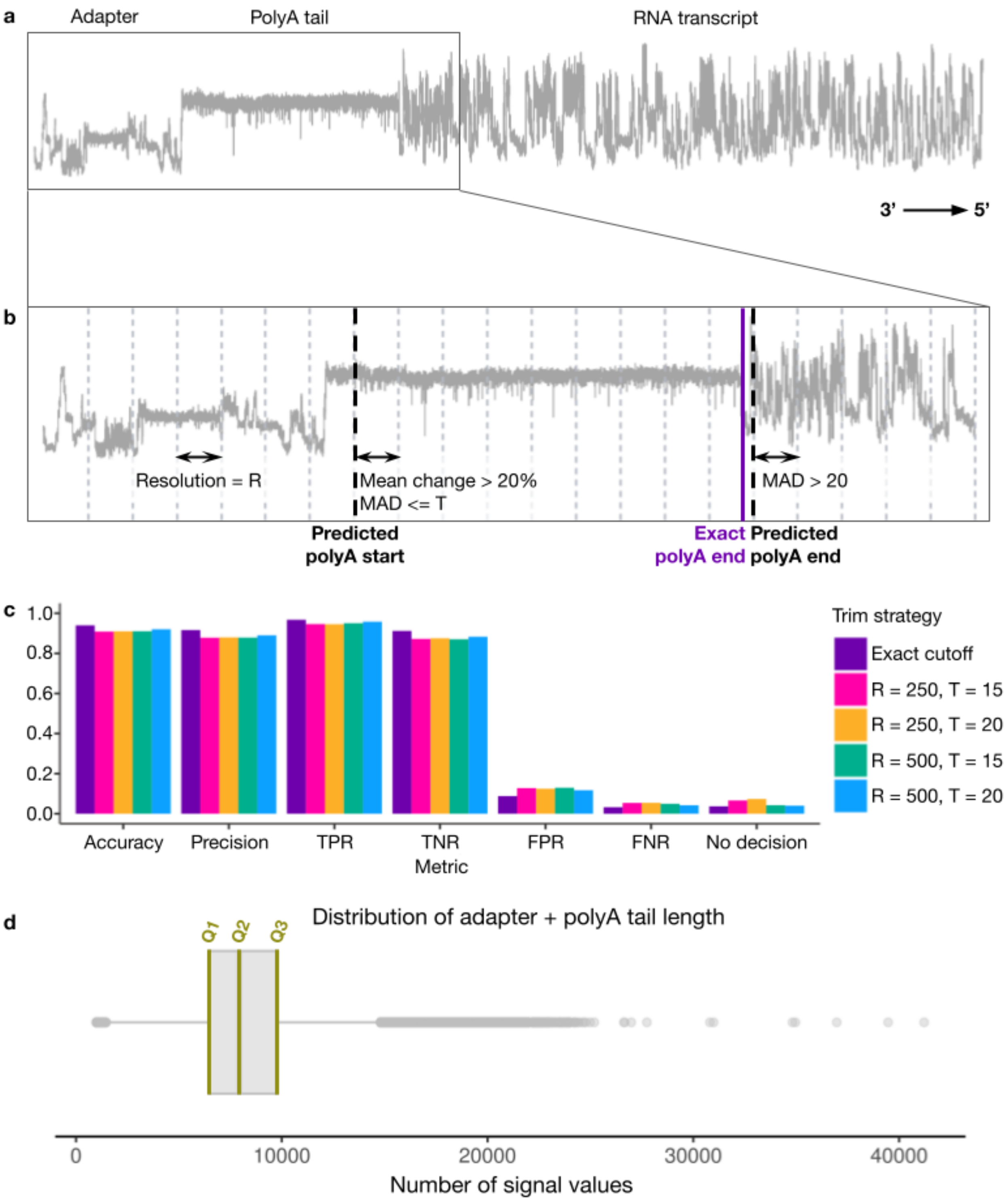


**Supplementary Figure 2: Removal of the sequencing adapter and poly(A) tail from the nanopore signal. a,** Example DRS signal from the 3' to 5' direction with adapter, poly(A) tail and RNA transcript portions of the signal annotated. **b,** 3' end of the signal shown in (a) with the predicted (black) and actual (purple) boundary between the poly(A) tail and RNA transcript shown, as well as the signal characteristics used for predicting the poly(A) tail position. **c,** Model performance on the mRNA model development test set to classify mRNA or non-mRNA when signals were trimmed at the exact boundary between the poly(A) tail and RNA transcript (purple), as well as by each candidate trimming strategy configuration: resolution (R) = 250 and initial median absolute deviation threshold (T) = 15 (pink), R = 250 and T = 20 (yellow), R = 500 and T = 15 (green) and R = 500 and T = 20 (blue). We show the metrics of accuracy, precision, true (TPR) and false

(FPR) positive rate, true (TNR) and false (FNR) negative rate and the proportion of signals for which a confident prediction could not be made. **d,** Distribution of the number of signal values (x-axis) comprising the adapter and poly(A) tail for the model development dataset, with quartiles highlighted.

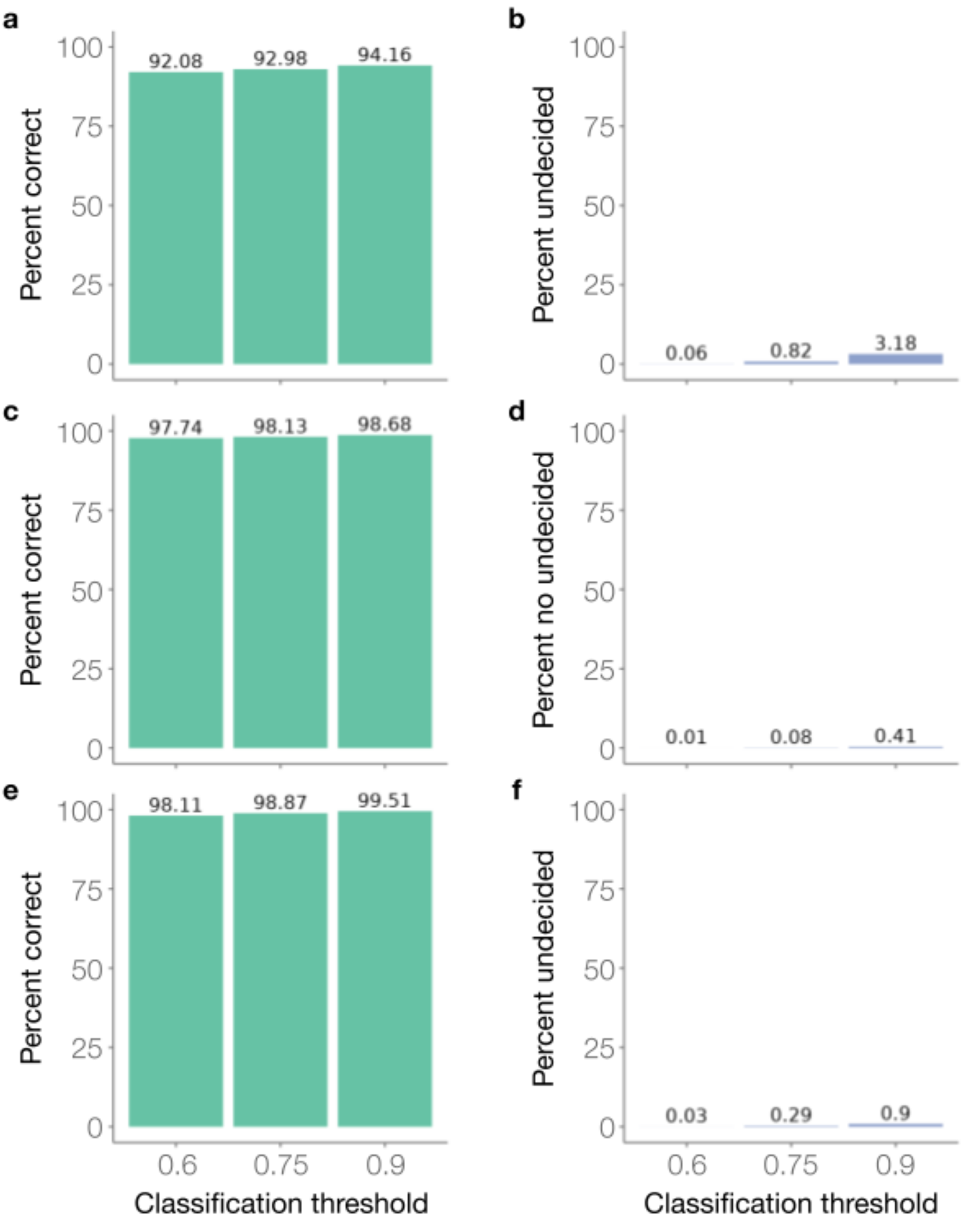


**Supplementary Figure 3: Classification threshold optimization.** For each candidate classification threshold (0.6: low, 0.75: medium and 0.9: high), we show the percentage of reads that RISER predicted correctly (**a**,**c**,**e**) or was unable to make a prediction for (**b**,**d**,**f**) in the respective test sets for the three RISER models implemented (mRNA: a-b, mtRNA: c-d, globin mRNA: e-f).

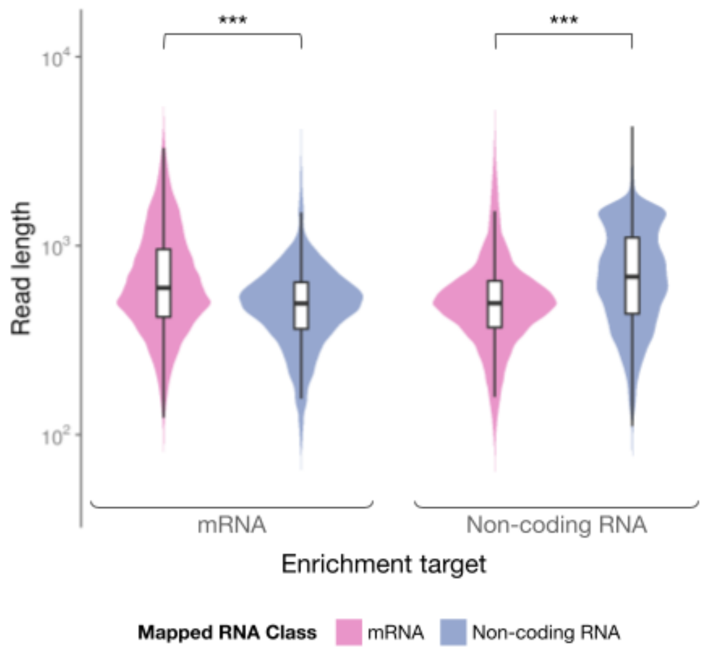


**Supplementary Figure 4: RISER reduces the length of off-target reads when used to enrich the target RNA class in a simulated live-sequencing environment.** The distribution of read lengths for the mapped RNA class when the RISER target class for enrichment was mRNA (left panel) and non-coding RNA (right panel) in a simulated live-sequencing run of an REH cancer cell line run using MinKNOW playback. The mapped RNA class is color-coded (pink: mRNA, blue: non-coding RNA). For both targets, the read lengths of the mapped RNA classes were compared using a Wilcoxon rank sum test with continuity correction ($H_1$: on-target > off-target) (p-value < 2.22E-16 for both target classes). Outliers were not included.

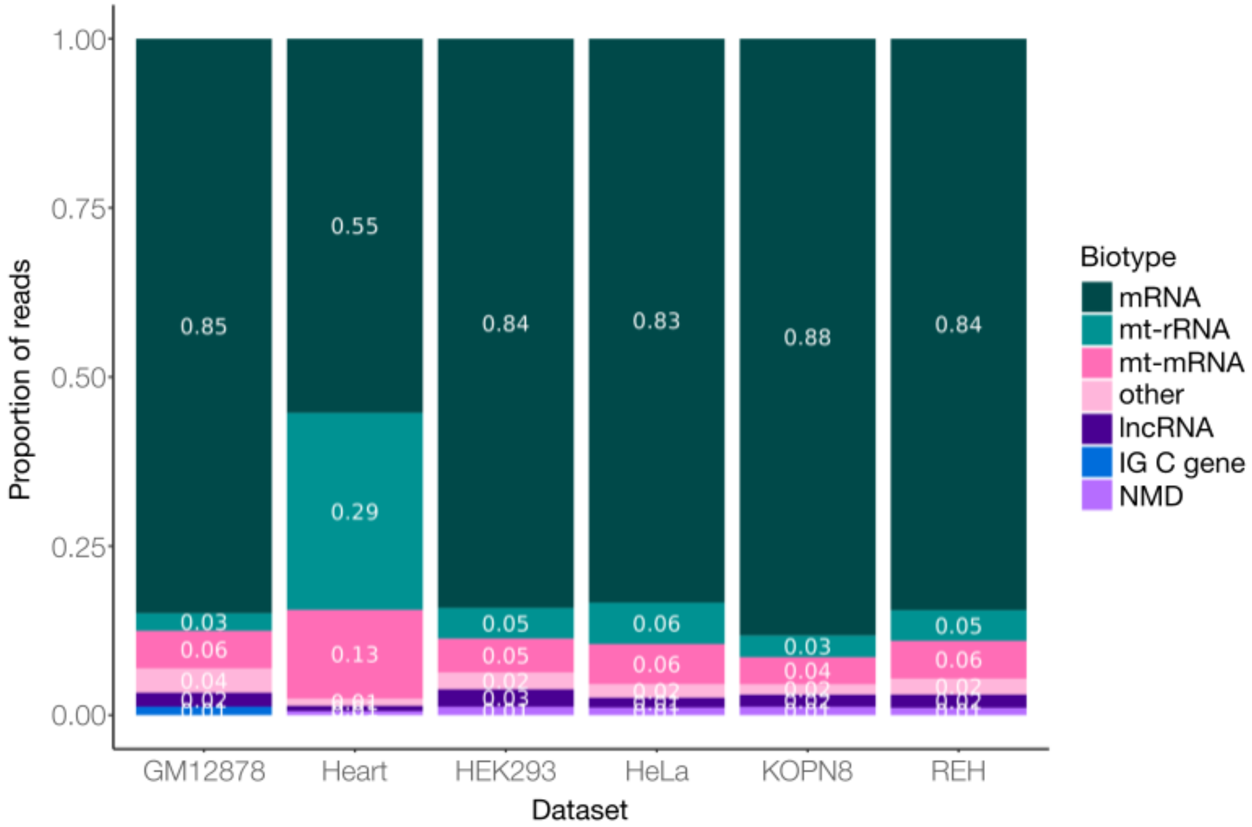


**Supplementary Figure 5: Proportion of reads per biotype in standard DRS runs across different cell lines.** The top 5 biotypes according to read proportion are displayed for each run, with the remaining biotypes aggregated under "other".

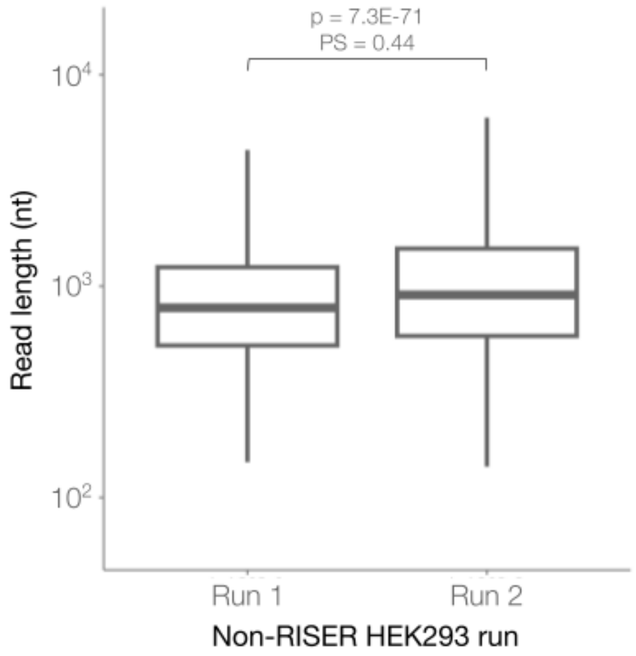


**Supplementary Figure 6: Read length distributions in non-RISER sequencing runs of HEK293 cells.** Distribution of read lengths (y-axis, log10-scale) for two MinION Mk1B sequencing runs performed without RISER. Outliers were not included. The read lengths of each run were compared using a Wilcoxon rank sum test (H1: run1 $\neq$ run2). The probability of superiority (PS) is also shown.

PS is the probability that a randomly sampled read from run 1 is longer than a randomly sampled read from run 2 (i.e., PS close to 0.5 means the lengths are likely to be the same, whereas PS close to 0 means that the run 1 lengths are highly likely to be shorter).

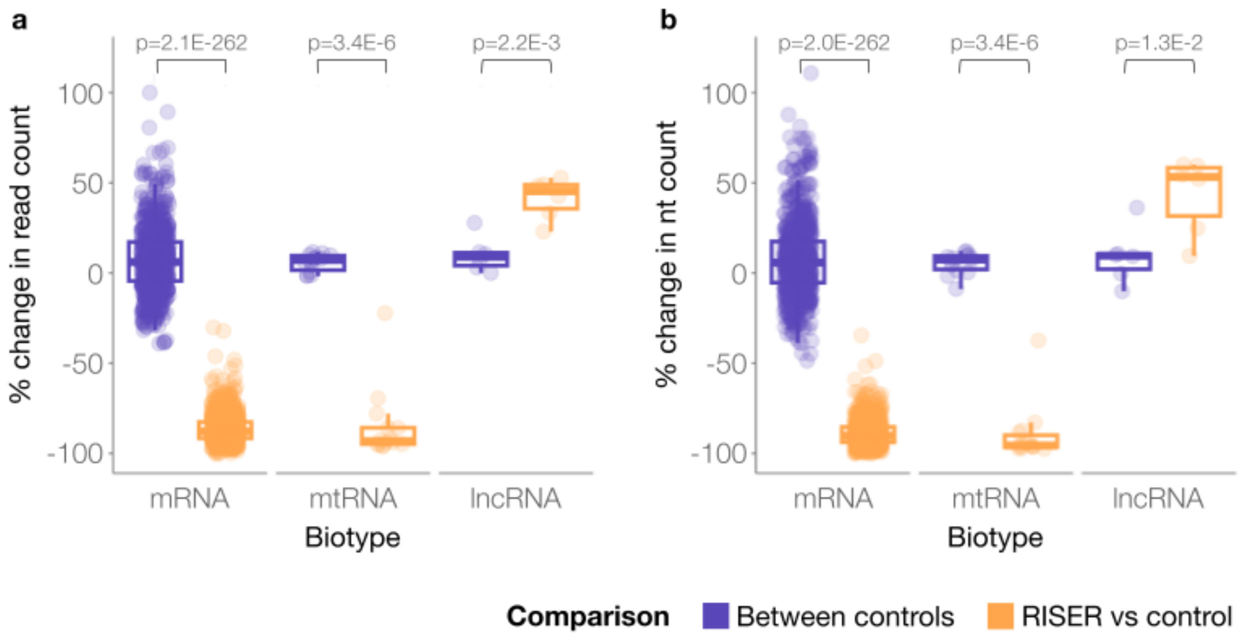


**Supplementary Figure 7: Effect of RISER on read and nucleotide counts.** Distribution of the percent change in read (**a**) and nucleotide (nt) (**b**) counts (y-axis) with respect to a control run, when RISER was used to deplete mRNA and mtRNA (orange) and for a separate control run (purple). For the set of transcripts in each biotype, the percent change in nt or reads using RISER was compared to the (no-RISER) control using a paired Wilcoxon signed rank test (H1 for mRNA and mtRNA: RISER<between controls, H1 for lncRNA: RISER>between controls). For this comparison, lncRNAs that did not overlap with coding exons from protein-coding transcripts were used.

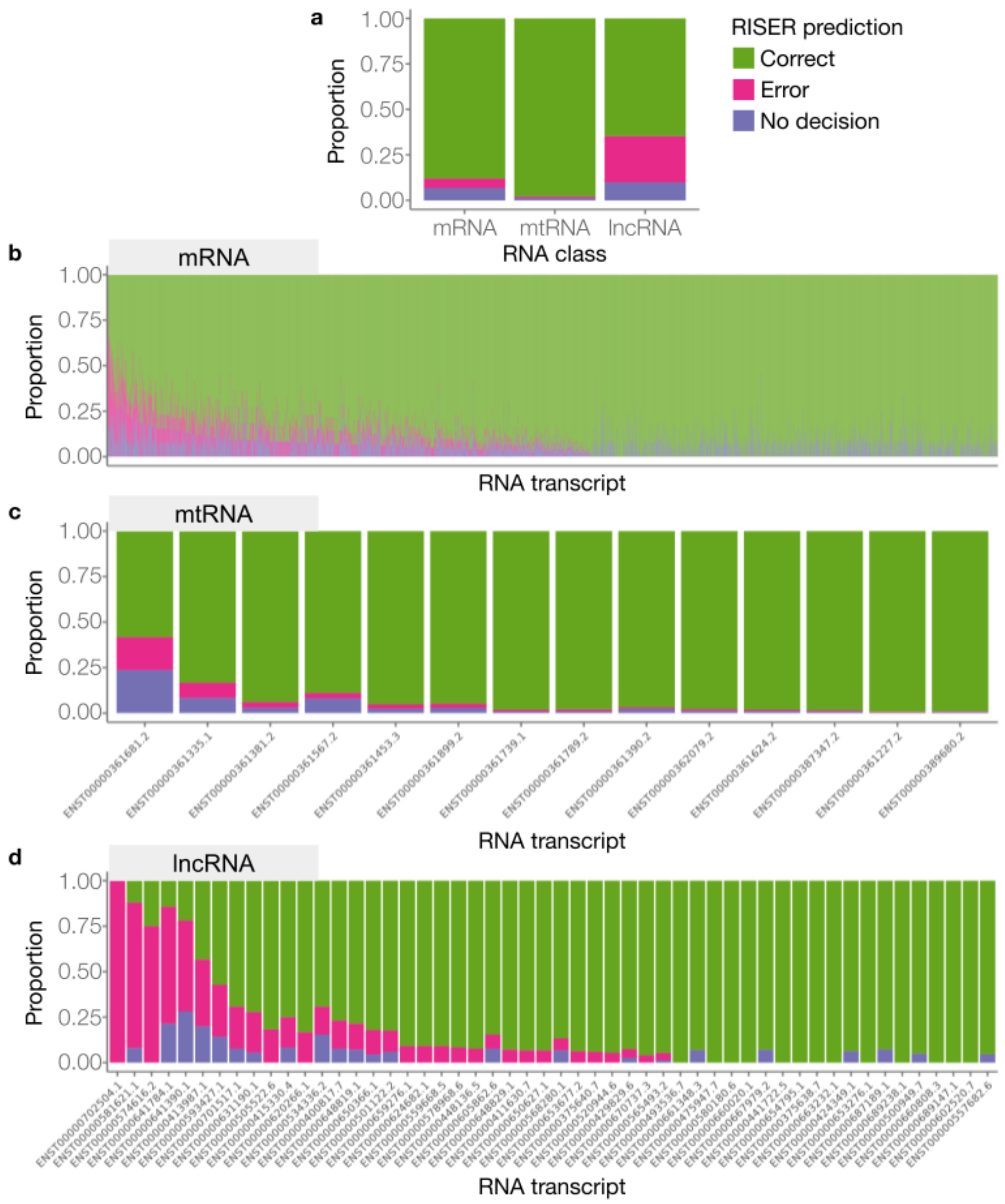


**Supplementary Figure 8. mRNA and mtRNA depletion transcript analysis.** We represent graphically the proportion of reads (y-axes) correctly classified (green), wrongly targeted by RISER (pink), or for which no decision was made (purple) during the live runs targeting mRNA and mtRNA for depletion. We represent these proportions for all mRNA, mtRNA and lncRNAs (**a**), and for all the individual transcripts from each class: mRNA (**b**), mtRNA (**c**) and lncRNA (**d**).

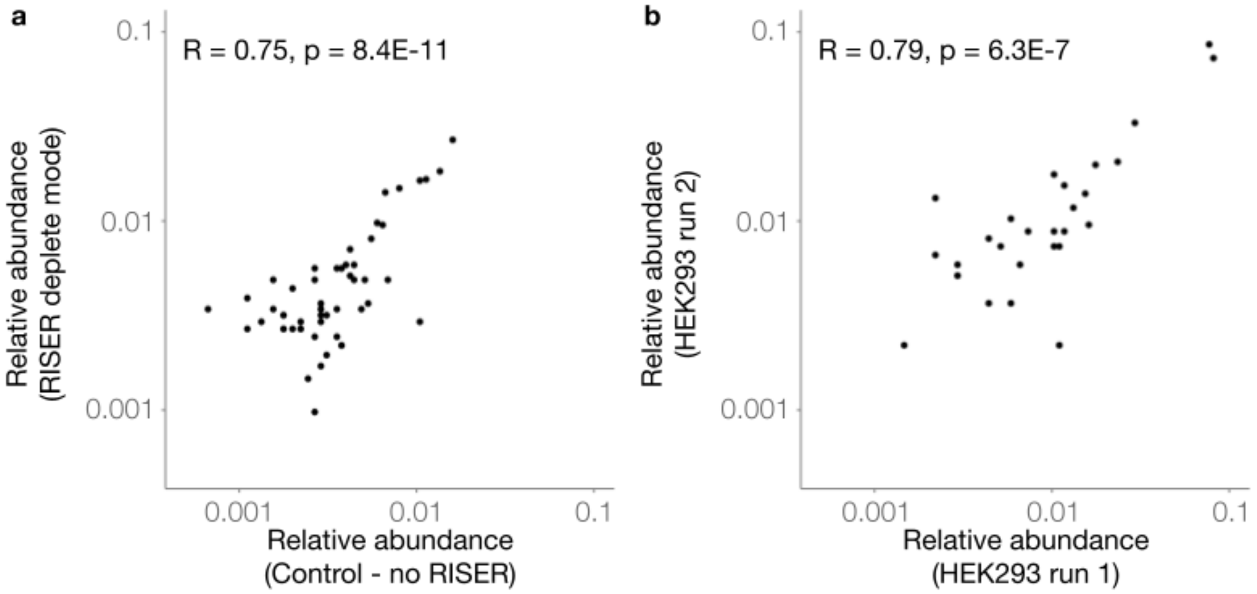


**Supplementary Figure 9: Conserved relative abundance of lncRNA after mRNA and mtRNA depletion. a,** For each lncRNA detected in the control (x-axis) and the RISER deplete mRNA and mtRNA (y-axis) conditions (in the same flow cell sequencing HEK293), we represent their abundance in log10 scale, measured as the proportion of reads mapped to that lncRNA over all reads mapped to all detected lncRNAs. Only lncRNAs with at least 10 reads in either the control or RISER condition were included. **b,** The abundance (calculated as in (a)) of lncRNAs measured in two separate sequencing experiments, without RISER, for the same cell line as in (a) (HEK293), considering the same number of reads as in (a), randomly sampled from the same set of transcripts as in (a). HEK293 DRS samples were prepared and sequenced with a MinION Mk1B as described in Methods.

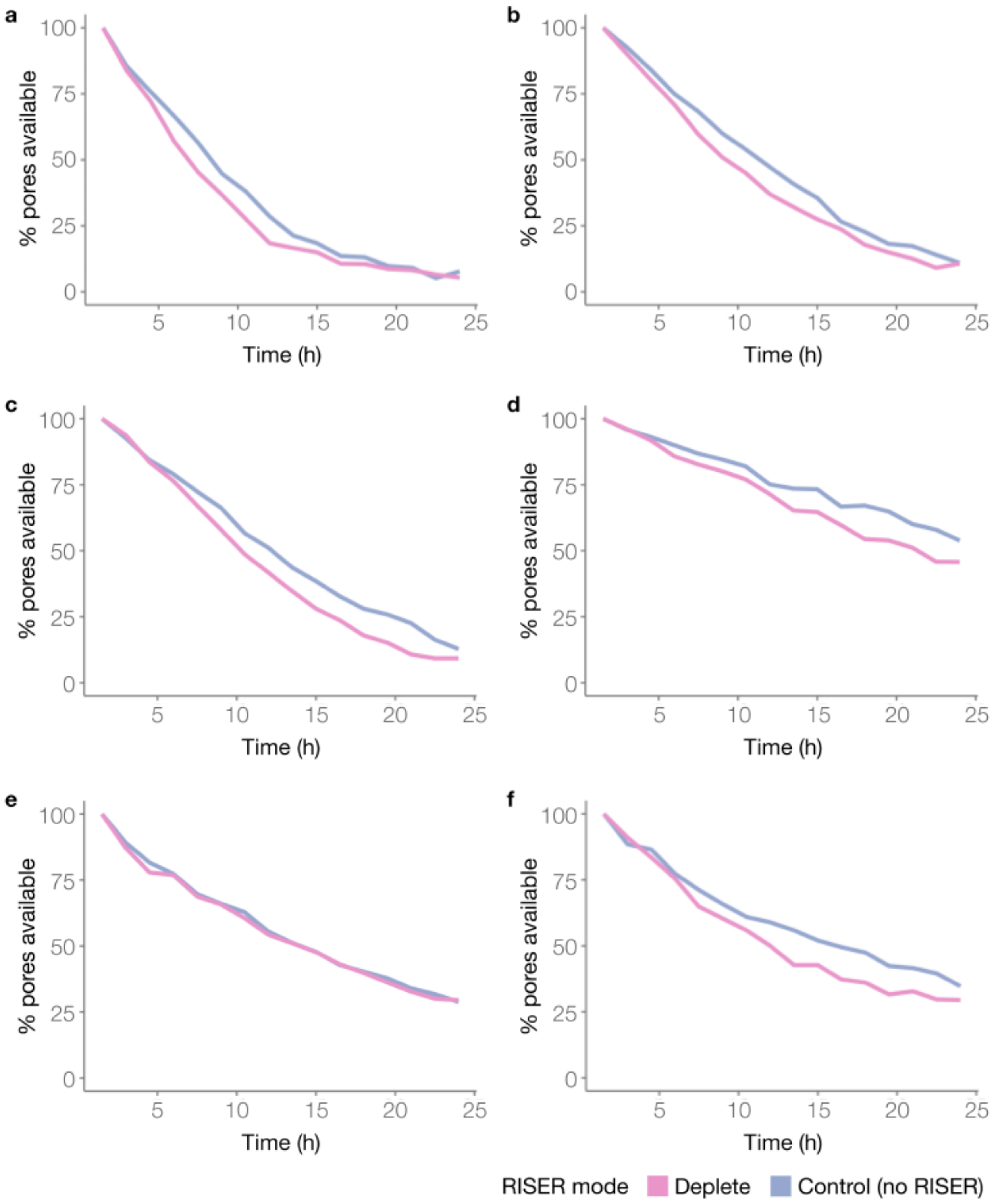


**Supplementary Figure 10: Percentage of available pores across 24h DRS runs.** We show the pores available over time as a percentage of the initial number of available pores for both RISER (pink) and control (blue) conditions, with RISER depleting mRNA and mtRNA in HEK293 (**a**, **b**) and HeLa (**c**) cell lines, and with RISER depleting globin mRNA in whole blood (**d-f**) samples. HEK293 DRS samples were prepared and sequenced as described in Methods. The HeLa sample was prepared as described in [1] and sequenced as described for HEK293 in Methods. The whole blood samples (**d-f**) were prepared and sequenced as described in Methods, except in (**f**), where total RNA from whole blood was extracted using the PAXgene blood RNA kit (Qiagen) as per manufacturer's instructions.

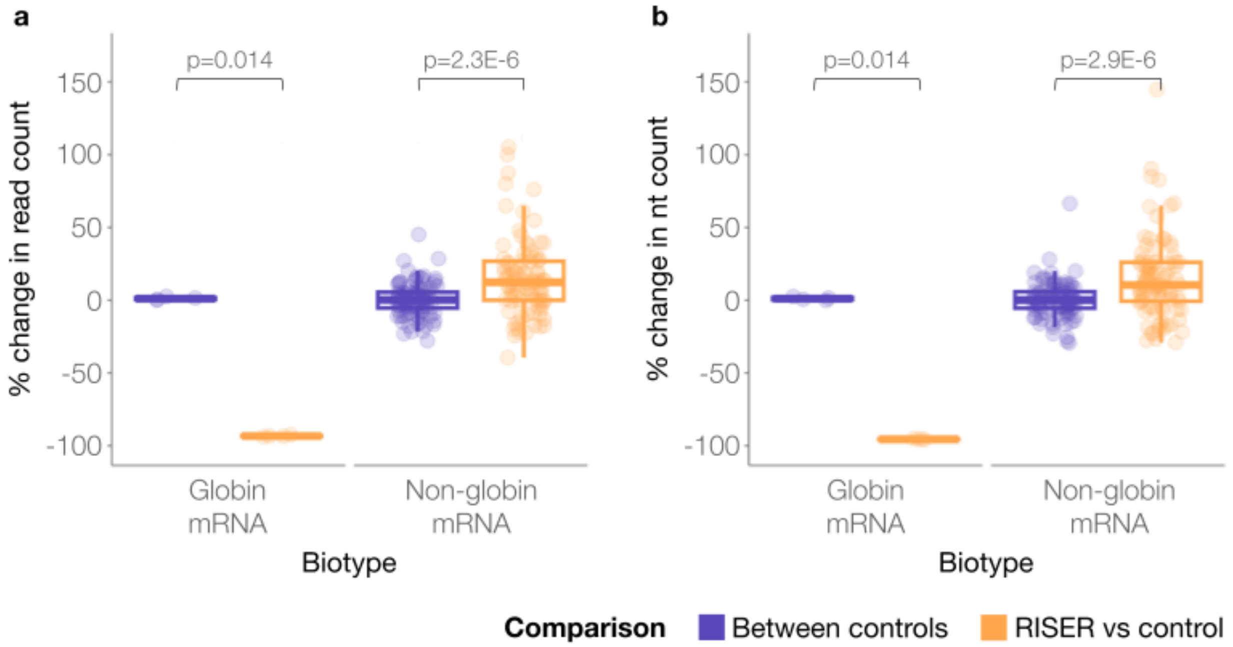


**Supplementary Figure 11: Effect of RISER on read and nucleotide counts.** Distribution of the percent change in read (**a**) and nucleotide (nt) (**b**) counts (y-axis), relative to a control run, when RISER was used to deplete globin mRNA (orange) and for a separate control run (purple). For the set of transcripts in each biotype, the percent change in nt or reads using RISER was compared to the (no-RISER) control using a paired Wilcoxon signed rank test (H1 for globin mRNA : RISER<between controls, H1 for non-globin mRNA: RISER>between controls).

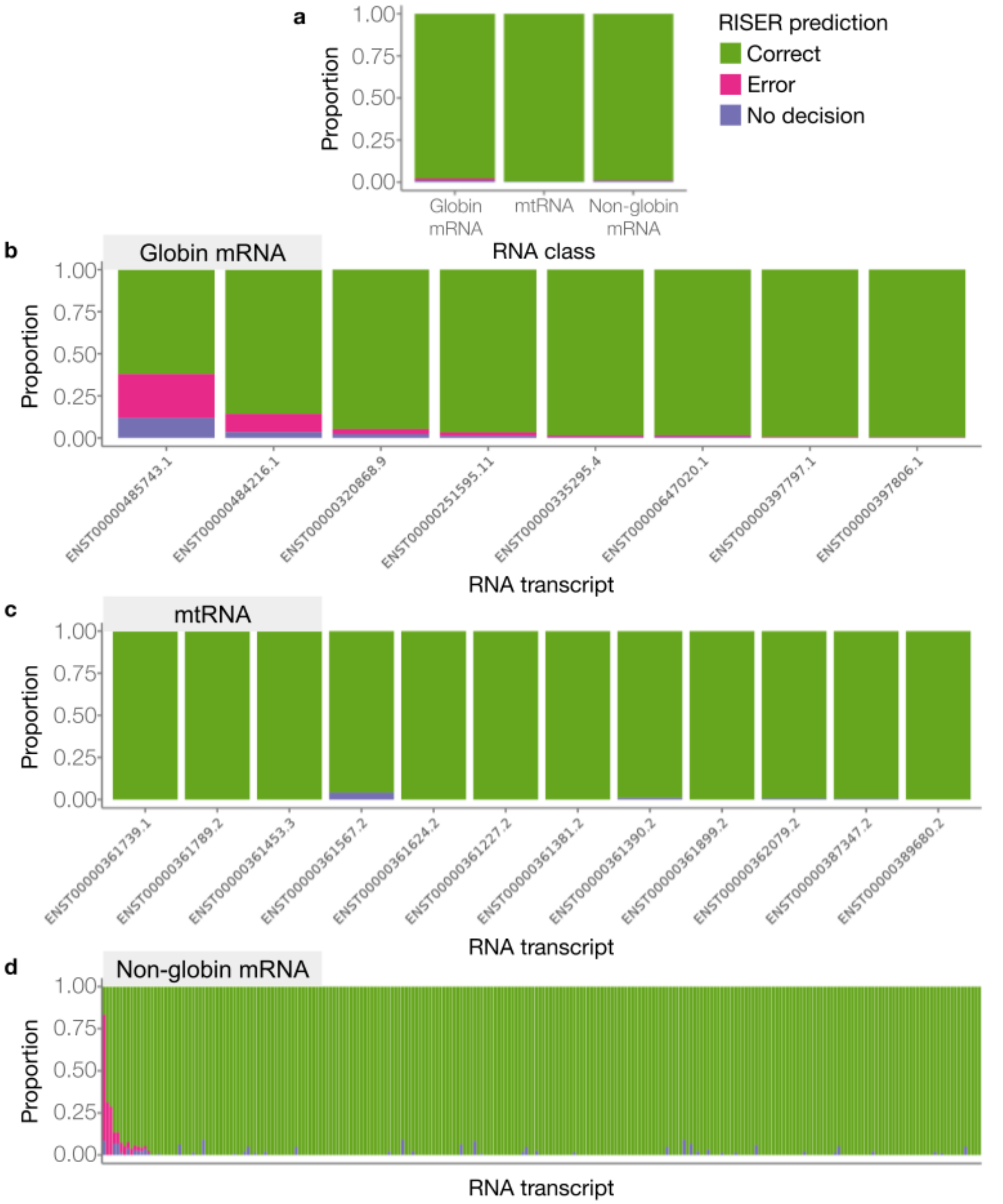


**Supplementary Figure 12: Globin depletion transcript analysis.** We represent graphically the proportion of reads (y-axes) correctly classified (green), wrongly targeted by RISER (fuchsia), or for which no decision was made (purple) during the live runs targeting globin mRNA for depletion. We represent these proportions for all globin mRNA, mtRNA and non-globin mRNA (**a**), and for all the individual transcripts from each class: globin mRNA (**b**), mtRNA (**c**) and non-globin mRNA (**d**).

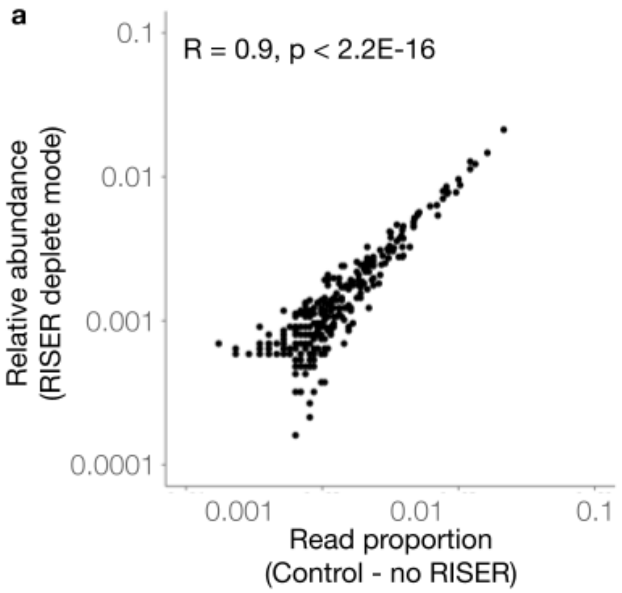


**Supplementary Figure 13: Conserved relative abundance of non-globin mRNA after globin depletion. a,** For each non-globin mRNA detected in the control (x-axis) and the RISER deplete globin mRNA (y-axis) conditions (in the same flow cell sequencing whole blood), we represent their abundance in log10 scale, measured as the proportion of reads mapped to that non-globin mRNA over all reads mapped to all detected non-globin mRNAs. Only non-globin mRNAs with at least 10 reads in either the control or RISER condition were included. **b,** The abundance (calculated as in (a)) of non-globin mRNAs measured in two separate sequencing experiments of whole blood without RISER, considering the same number of reads as in (a), randomly sampled from the same set of transcripts as in (a). Whole blood samples were prepared and sequenced with a MinION Mk1B as described for the production of globin model training data in Methods.

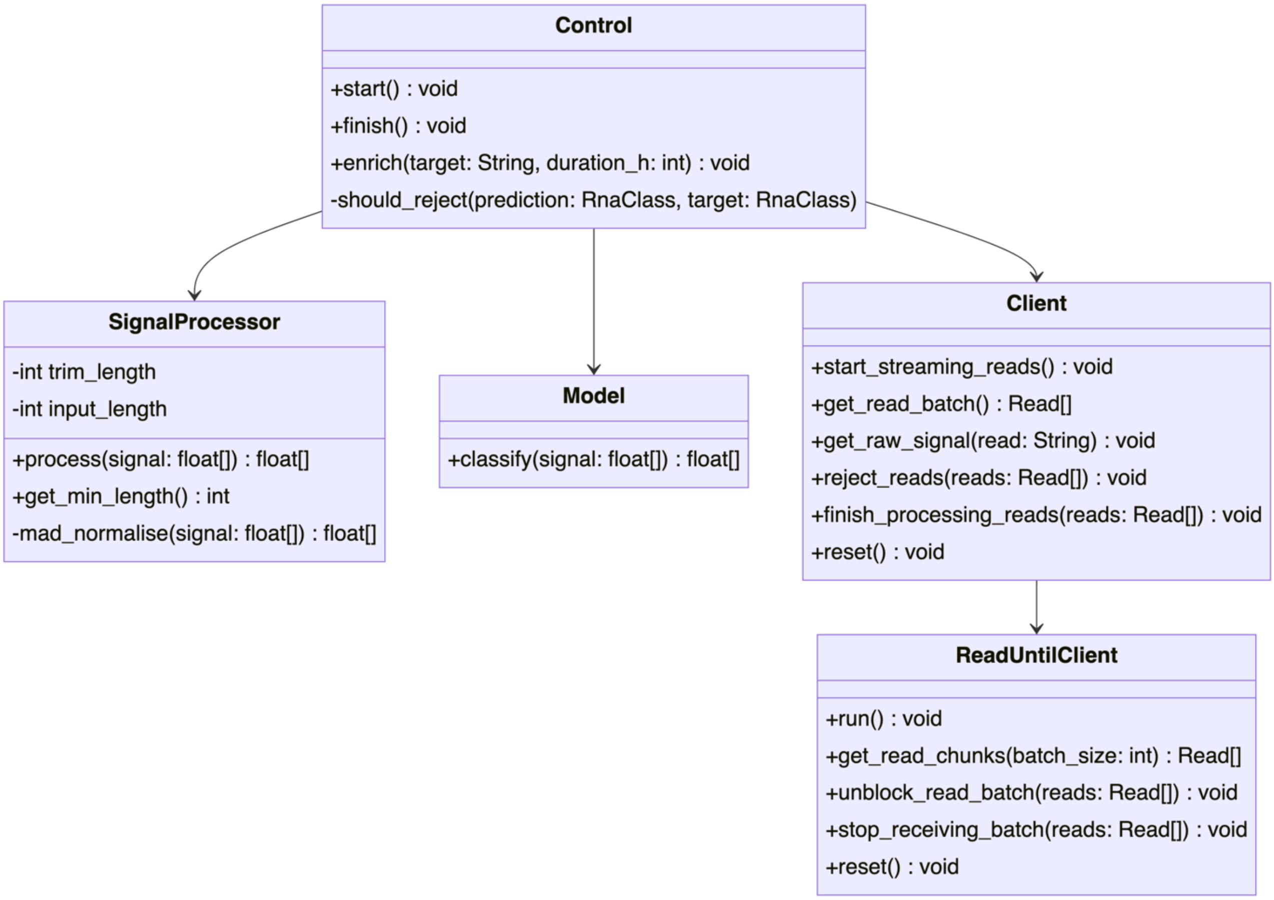


**Supplementary Figure 14: Class diagram for the RISER software using unified modelling language (UML) notation.** Each box represents a class and contains three sections: name, attributes, and method definitions. Attributes and method definitions have assigned access privileges, with ”+” and ”-" indicating public and private visibility, respectively. Arrows denote associations between classes, i.e., the class at the arrow's tail has an instance of the class at the tip of the arrow. For simplicity, logging, csv writing functionality and optional method parameters have not been shown. Only methods in the ONT ReadUntilClient (v3.0.0) that have been utilized by RISER are shown.

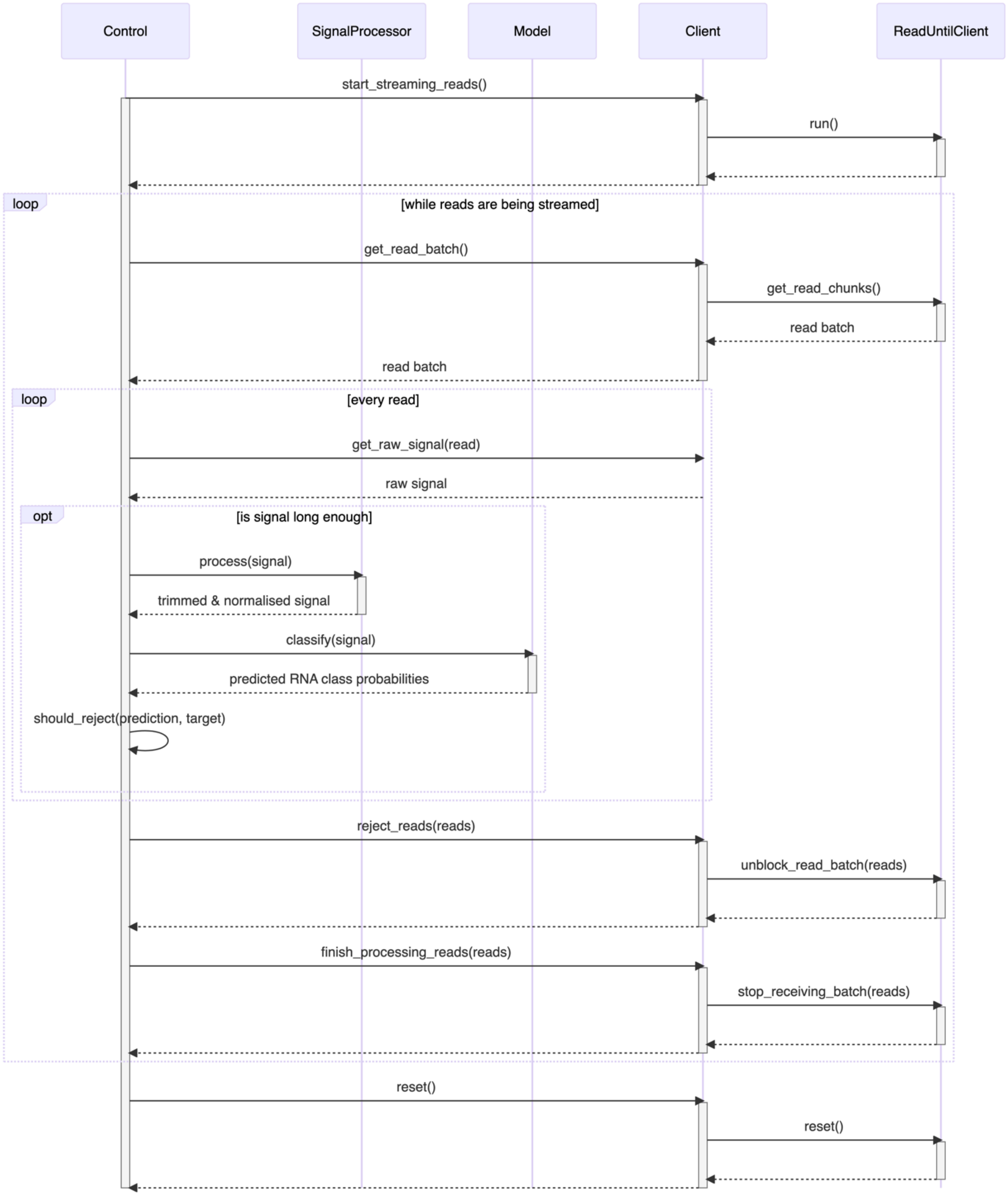


**Supplementary Figure 15: Sequence diagram for the RISER software using unified modelling language (UML) notation.** The diagram illustrates the timeline (top-to-bottom) of interactions between RISER components during a targeted sequencing run. Each blue box represents an object (class instance), with vertical blue lifelines extending beneath them. The white vertical bars denote when an object is busy (either executing a task or awaiting a message). Arrows denote synchronous messages (the sender requires a reply before continuing), with solid arrows representing messages from the sender to the receiver and dotted arrows denoting the return message from receiver to sender. Conditional functionality is enclosed within an "opt" rectangle, with the condition listed in square brackets. Loops are also enclosed within a "loop" rectangle, with the loop condition listed in square brackets.

#### Supplementary Tables

**Supplementary Table 1: MinION DRS datasets used in this study for the RISER model architecture development.** The HEK293-B dataset was composed by taking the second and third biological replicates for both the METTL3 knockout and wild-type samples in ENA PRJEB40872 and combining all technical replicates.

| Dataset ID | Cell line / tissue type | Accession # | Sequencing kit | Flow cell | Pore |
|---|---|---|---|---|---|
| GM24385 | GM24385 | TBD | SQK-RNA002 (modified) | FLO-MIN106 | R9.4.1 |
| Heart | Human heart | ENA PRJEB40410[2] | SQK-RNA002 | FLO-MIN106 | R9.4.1 |
| HEK293-A | HEK293 | TBD | SQK-RNA002 | FLO-MIN106 | R9.4.1 |
| HEK293-B | HEK293 | ENA PRJEB40872[3] | SQK-RNA002 | FLO-MIN106 | R9.4.1 |

**Supplementary Table 2: Hyperparameter optimization results for the Residual Network (ResNet) architecture.** "O" and "A" denote the "bottleneck" and "basic" residual block types, respectively, defined in [4].

| Input signal length | Batch size | Learning rate | Block type | Num. layers (l) | Kernel size (k) | Blocks (b) | Channels (c) | Num. trainable params | Val. accuracy (%) |
|---|---|---|---|---|---|---|---|---|---|
| 4s | 32 | 0.001 | O | 4 | 13 | 2,2,2,2 | 20,30,45,67 | 75,374 | 76.44 |
| 4s | 1000 | 0.001 | O | 4 | 13 | 2,2,2,2 | 20,30,45,67 | 75,374l | 75.26 |
| 4s | 32 | 0.001 | O | 4 | 19 | 2,2,2,2 | 16,24,36,54 | 49,086 | 76.23 |
| 4s | 16 | 0.001 | O | 4 | 19 | 2,2,2,2 | 20,30,45,67 | 75,494 | 76.67 |
| 4s | 32 | 0.0001 | O | 4 | 19 | 2,2,2,2 | 20,30,45,67 | 75,494 | 75.41 |
| 4s | 32 | 0.001 | O | 4 | 19 | 2,2,2,2 | 20,30,45,67 | 75,494 | 76.59 |
| 4s | 32 | 0.001 | O | 4 | 19 | 2,2,2,2 | 20,30,45,67 | 24,990 | 74.68 |
| 4s | 1000 | 0.0001 | O | 4 | 19 | 2,2,2,2 | 20,30,45,67 | 75,494 | 73.18 |
| 4s | 1000 | 0.001 | O | 4 | 19 | 2,2,2,2 | 20,30,45,67 | 75,494 | 75.41 |
| 4s | 32 | 0.001 | O | 4 | 19 | 2,2,2,2 | 20,40,80,160 | 278,602 | 76.8 |
| 4s | 32 | 0.001 | O | 4 | 19 | 2,2,2,2 | 32,48,72,108 | 191,562 | 76.75 |
| 4s | 32 | 0.001 | O | 4 | 19 | 2,2,2,2 | 32,64,128,256 | 708,418 | 77.08 |
| 4s | 1000 | 0.001 | O | 4 | 19 | 2,2,2,2 | 32,64,128,256 | 708,418 | 76.07 |
| 4s | 32 | 0.001 | O | 4 | 19 | 2,3,4,2 | 20,30,45,67 | 106,154 | 77.08 |
| 4s | 1000 | 0.001 | O | 4 | 19 | 2,3,4,2 | 20,30,45,67 | 106,154 | 75.92 |
| 4s | 32 | 0.001 | O | 4 | 19 | 3,4,23,3 | 20,30,45,67 | 379,514 | 78.75 |
| 4s | 32 | 0.001 | O | 4 | 19 | 3,4,6,3 | 128, 256, 512, 1024 | 7,670,402 | 77.73 |
| 4s | 32 | 0.001 | O | 4 | 19 | 3,4,6,3 | 20,30,45,67 | 166,844 | 78.1 |
| 4s | 32 | 0.001 | O | 4 | 19 | 3,4,6,3 | 256, 512, 1024, 2048 | 30,596,354 | 77.43 |

| | | | | | | | | | |
|---|---|---|---|---|---|---|---|---|---|
| 4s | 32 | 0.001 | O | 4 | 19 | 3,4,6,3 | 32,64,128,256 | 487,394 | 77.94 |
| 4s | 32 | 0.001 | O | 4 | 19 | 3,4,6,3 | 64, 128, 256, 512 | 1,928,258 | 78.16 |
| 4s | 32 | 0.001 | O | 4 | 25 | 2,2,2,2 | 20,30,45,67 | 75,614 | 76.56 |
| 4s | 1000 | 0.001 | O | 4 | 25 | 2,2,2,2 | 20,30,45,67 | 75,614 | 75.35 |
| 4s | 32 | 0.001 | O | 5 | 19 | 2,2,2,2,2 | 20,30,45,67,100 | 168,384 | 78.09 |
| 4s | 1000 | 0.001 | O | 5 | 19 | 2,2,2,2,2 | 20,30,45,67,100 | 168,384 | 76.8 |
| 4s | 32 | 0.001 | O | 6 | 19 | 2,2,2,2,2,2 | 20,30,45,67,100,150 | 375,684 | 79.32 |
| 4s | 32 | 0.001 | O | 7 | 19 | 2,2,2,2,2,2,2 | 20,30,45,67,100,150,225 | 840,384 | 80.16 |
| 4s | 32 | 0.001 | A | 4 | 19 | 2,2,2,2 | 20,30,45,67 | 102,120 | 78.23 |
| 4s | 1000 | 0.001 | A | 4 | 19 | 2,2,2,2 | 20,30,45,67 | 102,120 | 77.14 |
| 4s | 32 | 0.001 | A | 4 | 19 | 3,4,6,3 | 32,64,128,256 | 2,041,090 | 79.76 |
| 4s | 32 | 0.001 | A | 4 | 19 | 3,4,6,3 | 64, 128, 256, 512 | 8,139,266 | 79.36 |
| 4s | 32 | 0.0001 | A | 10 | 19 | 1,1,1,1,1,1,1,1,1,1 | 20,30,45,67,100,150,225,337,505,757 | 5,869,558 | 80.17 |
| 4s | **32** | **0.001** | **A** | **10** | **19** | **1,1,1,1,1,1,1,1,1,1** | **20,30,45,67,100,150,225,337,505,757** | **5,869,558** | **81.12** |

**Supplementary Table 3: Hyperparameter optimization results for the Temporal Convolutional Network (TCN) architecture.**

| Input signal length | Batch size | Learning Rate | Num. layers (l) | Num. channels (c) | Kernel size (k) | Dilation base (d) | Dropout (r) | # Trainable Params | % Val Accuracy |
|---|---|---|---|---|---|---|---|---|---|
| 4s | 32 | 0.0001 | 5 | 128 | 7 | 6 | 0.2 | 1,102,210 | 77.7 |
| 4s | 32 | 0.0001 | 5 | 128 | 7 | 8 | 0.2 | 1,102,210 | 76.07 |
| 4s | 32 | 0.0001 | 6 | 128 | 7 | 4 | 0.2 | 1,348,610 | 77.54 |
| 4s | 32 | 0.0001 | 7 | 128 | 7 | 3 | 0.2 | 1,595,010 | 76.97 |

| | | | | | | | | | |
|---|---|---|---|---|---|---|---|---|---|
| 4s | 32 | 0.0001 | 9 | 16 | 15 | 2 | 0.2 | 68,338 | 76.48 |
| 4s | 32 | 0.0001 | 10 | 16 | 7 | 2 | 0 | 37,314 | 76.35 |
| 4s | 32 | 0.0001 | 10 | 16 | 7 | 2 | 0.05 | 37,314 | 76.56 |
| 4s | 32 | 0.0001 | 10 | 16 | 7 | 2 | 0.1 | 37,314 | 76.29 |
| 4s | 16 | 0.0001 | 10 | 16 | 7 | 2 | 0.2 | 37,314 | 74.89 |
| 4s | 32 | 0.00001 | 10 | 16 | 7 | 2 | 0.2 | 37,314 | 72.46 |
| 4s | 32 | 0.0001 | 10 | 16 | 7 | 2 | 0.2 | 37,314 | 75.17 |
| 4s | 32 | 0.001 | 10 | 16 | 7 | 2 | 0.2 | 37,314 | 72.89 |
| 4s | 32 | 0.0001 | 10 | 16 | 9 | 2 | 0.2 | 47,074 | 76.03 |
| 4s | 32 | 0.0001 | 10 | 16 | 11 | 2 | 0.2 | 56,834 | 76.72 |
| 4s | 32 | 0.0001 | 10 | 32 | 7 | 2 | 0.2 | 147,330 | 76.49 |
| 4s | **32** | **0.0001** | **10** | **32** | **11** | **2** | **0.05** | **225,282** | **78.9** |
| 4s | 32 | 0.0001 | 10 | 64 | 7 | 2 | 0.2 | 585,474 | 77.37 |
| 4s | 32 | 0.0001 | 10 | 64 | 11 | 2 | 0.05 | 897,026 | 78.59 |
| 4s | 16 | 0.0001 | 10 | 128 | 7 | 2 | 0.2 | 2,334,210 | 76.7 |
| 4s | 32 | 0.0001 | 10 | 128 | 7 | 2 | 0.2 | 2,334,210 | 75.76 |
| 4s | 32 | 0.0001 | 11 | 16 | 5 | 2 | 0.2 | 30,450 | 73.68 |
| 4s | 32 | 0.0001 | 12 | 16 | 3 | 2 | 0.2 | 21,538 | 71.23 |
| 4s | 32 | 0.0001 | 12 | 128 | 3 | 2 | 0.2 | 1,319,170 | 75.59 |

**Supplementary Table 4: Hyperparameter optimization results for the “vanilla” Convolutional Neural Network (CNN) architecture.** The classifier “fc” denotes a 2-layer fully connected network with rectified linear unit (ReLu) activation, “gap” denotes a global average pooling layer and “gap_fc” denotes a global average pooling layer followed by a single fully connected layer.

| Input signal length | Batch size | Learning rate | Num. layers (l) | Blocks per layer (b) | Channels (c) | Kernel size (k) | Classifier (f) | # Trainable params | % Val accuracy |
|---|---|---|---|---|---|---|---|---|---|
| 4s | 32 | 0.001 | 4 | 1 | 20,30,45,67 | 3 | fc | 206,674,703 | 69.74 |
| 4s | 32 | 0.001 | 4 | 1 | 20,30,45,67 | 3 | gap | 15,253 | 70.08 |
| 4s | 32 | 0.001 | 4 | 1 | 20,30,45,67 | 3 | gap_fc | 15,523 | 70.15 |
| 4s | 32 | 0.001 | 4 | 1 | 20,30,45,67 | 5 | gap_fc | 25,223 | 72 |
| 4s | 32 | 0.001 | 4 | 1 | 20,30,45,67 | 7 | gap_fc | 35,193 | 72.08 |
| 4s | 32 | 0.001 | 4 | 1 | 20,30,45,67 | 11 | gap_fc | 55,133 | 73.45 |
| 4s | 32 | 0.001 | 4 | 1 | 32,64,128,256 | 3 | gap_fc | 130,114 | 70.79 |
| 4s | 32 | 0.001 | 4 | 2 | 20,30,45,67 | 3 | gap_fc | 38,857 | 72.05 |
| 4s | 32 | 0.001 | 4 | 4 | 20,30,45,67 | 3 | gap_fc | 86,065 | 50 |
| 4s | 32 | 0.0001 | 4 | 4 | 20,30,45,67 | 3 | gap_fc | 86,065 | 73.9 |
| 4s | 32 | 0.001 | 5 | 1 | 20,30,45,67,100 | 3 | gap_fc | 35,519 | 72.21 |
| 4s | 32 | 0.001 | 6 | 1 | 20,30,45,67,100,150 | 3 | gap_fc | 80,769 | 74.12 |
| 4s | 32 | 0.001 | 7 | 1 | 20,30,45,67,100,150,225 | 3 | gap_fc | 182,394 | 50 |
| 4s | 32 | 0.0001 | 7 | 1 | 20,30,45,67,100,150,225 | 3 | gap_fc | 182,394 | 76.51 |
| 4s | 32 | 0.001 | 8 | 1 | 20,30,45,67,100,150,225,337 | 3 | gap_fc | 410,430 | 50 |
| 4s | 32 | 0.0001 | 8 | 1 | 20,30,45,67,100,150,225,337 | 3 | gap_fc | 2,069,942 | 79.38 |
| 4s | 32 | 0.0001 | 8 | 1 | 20,40,80,160,320,640,1280,2560 | 3 | gap_fc | 13,116,682 | 78.99 |
| 4s | 32 | 0.0001 | 8 | 1 | 32,64,128,256,512,1024,2048,4096 | 3 | gap_fc | 33,568,834 | 79.29 |
| 4s | 32 | 0.001 | 10 | 1 | 20,20,30,30,45,45,67,67,100,100 | 3 | gap_fc | 89,223 | 50 |
| 4s | 32 | 0.001 | 10 | 1 | 20,30,45,67,100,150,225,337,505,757 | 3 | gap_fc | 2,069,942 | 50 |
| 4s | 32 | 0.0001 | 10 | 1 | 20,30,45,67,100,150,225,337,505,757 | 3 | gap_fc | 2,069,942 | 79.35 |

| | | | | | | | | | |
|---|---|---|---|---|---|---|---|---|---|
| 4s | 32 | 0.0001 | 10 | 2 | 20,30,45,67,100,150,225,337,505,757 | 3 | gap_fc | 5,169,924 | 50 |
| 4s | 32 | 0.00001 | 10 | 2 | 20,30,45,67,100,150,225,337,505,757 | 3 | gap_fc | 5,169,924 | 50 |
| 4s | 32 | 0.000001 | 10 | 2 | 20,30,45,67,100,150,225,337,505,757 | 3 | gap_fc | 5,169,924 | 50 |
| 4s | 32 | 0.0000001 | 10 | 2 | 20,30,45,67,100,150,225,337,505,757 | 3 | gap_fc | 5,169,924 | 50 |
| 4s | **32** | **0.0001** | **12** | **1** | **20,30,45,67,100,150,225,337,505,757,1135,1702** | **3** | **gap_fc** | **10,447,564** | **80.22** |

**Supplementary Table 5: Time taken to classify 1000 fixed-length DRS signals by sequence-based adaptive sampling (AS) and by RISER.** AS was performed using basecalling and mapping to the protein-coding transcriptome.

| Approach | Average classification time per signal (using GPU) (s) | Average classification time per signal (using CPU) (s) |
|---|---|---|
| RISER | 0.00585 | 0.0157 |
| AS | 0.258 | H5.83 |

**Supplementary Table 6: MinION DRS datasets used in this study for development and evaluation of the mRNA and mtRNA models.** The HEK293-B dataset was composed by taking the second and third biological replicates for both the METTL3 knockout and wild-type samples in ENA PRJEB40872 and combining all technical replicates. Only data from HeLa WT cells was used from GSE211762. (1) All JHU runs 1-5 and (2) Bham run 1 from the Nanopore WGS consortium https://github.com/nanopore-wgs-consortium/NA12878/

| Dataset ID | Cell line / tissue type | Accession # | Sequencing kit | Flow cell | Pore | Used for training? |
|---|---|---|---|---|---|---|
| GM12878-A | GM12878 | (1) | SQK-RNA001 | FLO-MIN106 | R9.4.1 | Yes |
| GM12878-B | GM12878 | TBD | SQK-RNA002 | FLO-MIN106 | R9.4.1 | Yes |
| GM12878-C | GM12878 | (2) | SQK-RNA001 | FLO-MIN106 | R9.4.1 | No |
| GM24385 | GM24385 | (1) | SQK-RNA002 (modified) | FLO-MIN106 | R9.4.1 | Yes |
| Heart | Human heart | ENA PRJEB404102 | SQK-RNA002 | FLO-MIN106 | R9.4.1 | Yes |
| HEK293-A | HEK293 | TBD | SQK-RNA002 | FLO-MIN106 | R9.4.1 | Yes |
| HEK293-B | HEK293 | ENA PRJEB408724 | SQK-RNA002 | FLO-MIN106 | R9.4.1 | Yes |
| HEK293-C | HEK293 | TBD | SQK-RNA002 | FLO-MIN106 | R9.4.1 | Yes |
| HeLa | HeLa | TBD | SQK-RNA002 | FLO-MIN106 | R9.4.1 | No |
| KOPN8 | KOPN8 | TBD | SQK-RNA002 | FLO-MIN106 | R9.4.1 | Yes |

| | | | | | | |
|---|---|---|---|---|---|---|
| REH | REH | TBD | SQK-RNA002 | FLO-MIN106 | R9.4.1 | Yes |

**Supplementary Table 7: RISER output files - CSV file.**

| read_id | channel | probability_noncoding | probability_coding | prediction | target | decision |
|---|---|---|---|---|---|---|
| 075391a9-2816-45b0-aebb-12b1f398fcd3 | 204 | 0.83 | 0.17 | NONCODING | CODING | REJECT |
| afcbd456-1843-4322-85d2-7f001ef0dc01 | 176 | 0.24 | 0.76 | CODING | CODING | ACCEPT |
| d8de6be4-4a01-42dc-b8ab-77abc8f818e1 | 373 | 0.36 | 0.64 | CODING | CODING | ACCEPT |
| 02cdeae6-3d5d-4615-bc61-7d5dd9a7217c | 91 | 0.79 | 0.21 | NONCODING | CODING | REJECT |
| 4460c783-4663-4666-be4d-c52590fdff31 | 293 | 0.26 | 0.74 | CODING | CODING | ACCEPT |
| … | … | … | … | … | … | … |

#### Supplementary Notes

##### Supplementary Note 1: RISER command structure.

```
usage: riser.py [-h] -t -m -d [--min] [--max] [--threshold]


optional arguments:
 -h, --help          show this help message and exit
 -t, --target        RNA class to enrich for. This must be one or more of
                     {mRNA,mtRNA,globin}. (required)
 -m, --mode          Whether to enrich or deplete the target class. This
                     must be one of {enrich,deplete} (required)
 -d, --duration      Length of time (in hours) to run RISER for. This
                     should be the same as the MinKNOW run length.
                     (required)
 --min               Minimum number of seconds of transcript signal to use
                     for RISER prediction. (default: 2)
 --max               Maximum number of seconds of transcript signal to try
                     to classify before skipping this read. (default: 4)
 --threshold         Probability threshold for classifier [0,1]. (default:
                     0.9)
```

##### Supplementary Note 2: RISER output files - Console output.

```
Using cuda device
Usage: riser.py -t noncoding -d 48
All settings used (including those set by default):
--target        : Class.NONCODING
--duration_h    : 48
--config_file   : models/cnn_best_model.yaml
--model_file    : models/cnn_best_model.pth
--polyA_length  : 6481
--secs          : 4
Client is running.
Batch of 110 reads received: 59 long enough to assess, 46 of which were
rejected (took 0.3148s)
Batch of  93 reads received: 29 long enough to assess, 21 of which were
rejected (took 0.1376s)
Batch of 107 reads received: 32 long enough to assess, 24 of which were
rejected (took 0.1568s)
...
```

##### Supplementary Note 3: RISER output files - Log file.

```
2022-08-23T11:56:29 [RISER] INFO: Using cuda device
2022-08-23T11:56:31 [RISER] INFO: Usage: riser.py --target noncoding --
duration 24
2022-08-23T11:56:31 [RISER] INFO: All settings used (including those set
by default):
2022-08-23T11:56:31 [RISER] INFO: --target        : Class.NONCODING
2022-08-23T11:56:31 [RISER] INFO: --duration_h    : 24
```

```
2022-08-23T11:56:31 [RISER] INFO: --config_file   :
defaults/cnn_best_model.yaml
2022-08-23T11:56:31 [RISER] INFO: --model_file    :
defaults/cnn_best_model.pth
2022-08-23T11:56:31 [RISER] INFO: --polyA_length  : 6481
2022-08-23T11:56:31 [RISER] INFO: --secs          : 4
2022-08-23T11:56:31 [RISER] INFO: Client is running.
2022-08-23T11:56:32 [RISER] INFO: Batch of   0 reads received:  0 long
enough to assess,  0 of which were rejected (took 0.0000s)
2022-08-23T11:56:33 [RISER] INFO: Batch of 268 reads received:  0 long
enough to assess,  0 of which were rejected (took 0.0013s)
2022-08-23T11:56:34 [RISER] INFO: Batch of 284 reads received:  0 long
enough to assess,  0 of which were rejected (took 0.0014s)
2022-08-23T11:56:35 [RISER] INFO: Batch of 262 reads received:  0 long
enough to assess,  0 of which were rejected (took 0.0015s)
...
```
